# Calibration and Evaluation of a Force Measurement Glove for Field-Based Monitoring of Manual Wheelchair Users

**DOI:** 10.1101/2020.07.19.169862

**Authors:** Anthony Anderson, Alexander W. Hooke, Chandrasekaran Jayaraman, Adam Burns, Emma Fortune, Jacob J. Sosnoff, Melissa M. Morrow

**Affiliations:** *Dept. of Mechanical Engineering, University of Washington*, Seattle, WA, USA; *Biomechanics Core Facility, Mayo Clinic*, Rochester, MN, USA; *Dept. of Kinesiology and Comm. Health, University of Illinois at Urbana-Champaign*, Urbana-Champaign, IL; *Div. of Health Care Policy and Research, Dept. of Health Sciences Research, Mayo Clinic*, Rochester, MN, USA

**Keywords:** wheelchair, spinal cord injury, force glove, kinetics

## Abstract

Manual wheelchair users exhibit pain related to repetitive and demanding shoulder activities of daily living. Wearable sensing systems like force measurement gloves can provide insights about upper extremity loading in the community and home environments. We calibrated and evaluated the accuracy of a novel force measurement glove with a body-worn data logger. The device was calibrated with loads of 0-800 N applied to the palmar surface of the glove. Calibration conditions were tested that varied the stiffness of the material in the glove, the temperature, and the curvature of the force applicator. Calibration equations from each condition were evaluated by comparing the glove’s force prediction with the output of an instrumented wheelchair rim during propulsion and weight relief exercises. The force measurement glove detected 72.7% of 355 propulsion peaks and had a strong linear correlation with the instrumented rim force measurements (r=0.80). The most accurate calibration equation was constructed using data from all conditions, with an RMS force measurement error of 64.7 N and 31.7 N for weight relief exercise and propulsion, respectively. The force measurement glove design described here may serve as a useful tool for detection of loading events and relative magnitude changes.

## I. Introduction

Manual wheelchair users suffer from high rates of upper extremity pain and injury [1]. Following a spinal cord injury that leads to use of a manual wheelchair for mobility, the upper extremities become responsible for independent locomotion. Weight relief exercises and other high-load events such as body transfers become activities of daily living. This increase in shoulder loading has the potential to cause subacromial bursitis, rotator cuff injuries, and other painful musculotendinous pathologies. [2].

While upper extremity loading in manual wheelchair users can be analyzed in a laboratory environment, studies have shown that clinically relevant biomechanical variables differ significantly in community settings, including average propulsion moment and mechanical work [3], [4]. However, field-based data collections require compromise as wearable sensors have drawbacks in comparison to laboratory-based sensing technologies. In a laboratory setting, shoulder loading during wheelchair propulsion can be estimated with optical motion capture systems, instrumented wheelchair rims, and a standard inverse dynamics approach [5]. While upper extremity orientation can be estimated with inertial measurement units in the field [6], technologies for the measurement of hand contact forces and moments in the field for full day data collections are limited. Measurement of forces during daily life is needed to understand joint loading contributions to overuse injury.

Instrumented wheels [3] can provide high-resolution biomechanical load measurements in the field; however, there are cost and feasibility limitations for using the instrumented wheels for full day and multi-day collections. other commercial force sensors tend to be mechanically rigid and are optimized for highly controlled loading scenarios. Estimating shoulder loads during wheelchair use requires measuring contact forces at the palms, which are highly variable as loading surfaces and require sensing modalities that can flex and conform as hand pose changes. To our knowledge, there is currently no sensing system capable of reliably measuring hand contact forces for extended data collections in the field.

Without access to high quality wearable force sensors, biomechanists have used force sensitive resistors (FSRs) to measure body contact forces in a variety of applications [7]. FSRs are thin-film sensors with electrical impedances that vary with mechanical load. FSRs are appealing to scientists because they are low-profile, flexible, and inexpensive. However, it is well known that FSR outputs are sensitive to temperature, curvature, loading area, and substrate stiffness [8], making them unsuitable for many biomechanical applications where loading conditions are highly variable. Recently, several research groups have investigated advanced calibration techniques that account for changes in the biomechanical variables that are most likely to influence FSR output [8–10].

In this paper, we describe the development and evaluation of a low-fidelity force measurement glove for extended field-based data collections with manual wheelchair users. This glove was used in one previous study to measure hand contact forces while using a forearm support walker [11]. While the glove accuracy was deemed adequate in that loading scenario, it was unknown if the FSRs would be able to reliably measure hand contact forces in the relatively unconstrained loading scenarios of manual wheelchair propulsion and weight relief exercises.

The purpose of this study was to evaluate the novel force measurement glove and a biomechanically informed calibration technique by comparing the calibrated glove’s predicted force to forces measured by an instrumented wheelchair rim during propulsion and weight relief exercises.

## II. Methods

### A. Glove Description

The glove consists of a cloth glove with four FSRs (Alibaba, Hangzhou, China) in the palm (Fig. 1). Each FSR has a fitted cloth housing that is sewn into the palm of the glove. The FSRs are 38 mm in diameter, and each sensor slightly overlaps with the neighboring sensors.

**Fig. 1:**
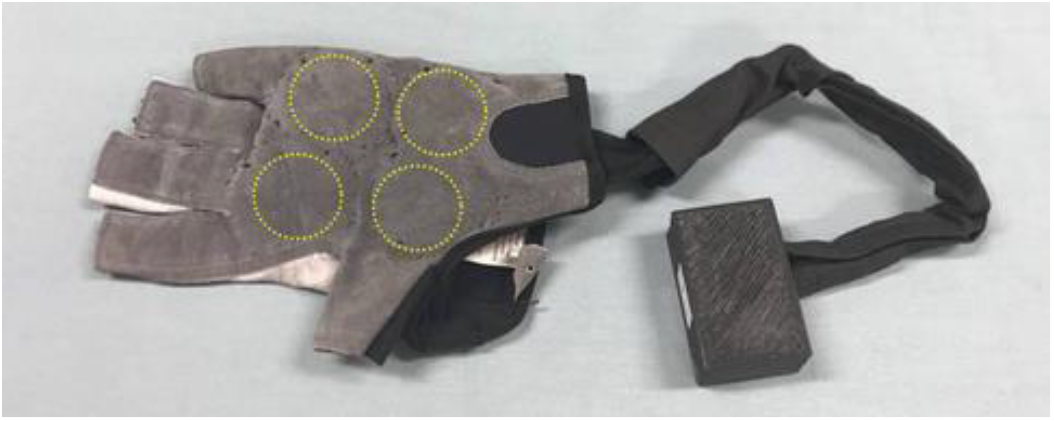
Force-sensing glove with custom printed circuit. Location of the forcesensing resistors are outlined in yellow circles.

The FSRs are connected to a small printed circuit board (PCB) through a protective cloth sleeve. The PCB sits inside of a 3D printed case that is fixed to the user’s forearm with an elastic band. The circuit is powered by a 3.7 V, 3000 mAh lithium polymer battery that sits inside of the 3D printed case during use. The PCB samples the sensor data and logs it to a 32 Gb microSD card. Sensor signals are sampled at 20 Hz with a 10-bit analog-to-digital converter. Data can be collected for several hours per charge.

### B. Calibration Testing

During the calibration protocol, known loads were applied to the palm of the glove in five conditions (Table I). For each condition, an MTS 585 Bionix II material testing system (MTS, Eden Prairie, MN) was used to apply loads between 0 N and 800 N to the surface of the palm of the glove in 80 N increments. This range was selected to encompass the expected static loading on one hand if subject weighing 1600 N were to support half of their body weight on one hand. A load cell attached in series with a force applicator provided feedback to the MTS control system. MTS force was held constant for ten seconds at each load and the glove was unloaded between loading conditions. The loading procedure was completed for five conditions based on findings from [8], with variations in sensor substrate stiffness, glove temperature, the curvature of the force applicator, or some combination of these variables. These specific loading conditions were selected to approximate the range of substrate stiffnesses, curvatures, and temperatures at the sensor-hand interface during manual wheelchair use.

Three different stiffnesses were tested during calibration. The stiffest condition resulted from loading the glove with a flat force applicator while it sat on a metal base plate within the MTS. The other two stiffnesses were achieved by inserting rectangular pieces of ballistic gel or artificial flesh (Syndaver, Miami, FL) inside the glove to simulate human soft-tissue compliance.

For condition four, the glove’s interior temperature was held at 35 C with a flexible heating pad that was wrapped around the artificial flesh material. For condition five, the glove was loaded with a custom force applicator made from a section of a wheelchair rim rather than the flat force applicator.

Matlab’s *polyfit* command (Matlab 2018b, The MathWorks, Natick, MA) was used to fit a linear equation to the calibration data for each condition. The equation (1) maps the sum of the four sensors’ ADC output (proportional to voltage) to the applied MTS load:

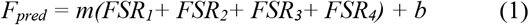

where Fpred is the force prediction of the glove, FSRi is the bit reading of the i’th FSR, and m and b are calibration coefficients. FSR readings were summed due to the overlap of the sensors. Pilot testing showed no improvement in force prediction when each sensor was assigned a unique gain. Another equation was fit using data from all conditions and was evaluated as a sixth condition. In all cases, the glove force predictions were modified to predict no load when the sensor output was zero.

### C. Glove Force Prediction in Comparison to SmartWheel

A procedure was developed to compare the glove force predictions for each calibration condition with the output of a SmartWheel (Three Rivers Holdings Inc., Mesa, AZ) during wheelchair propulsion and weight relief exercises. A SmartWheel is an instrumented wheelchair rim that measures three-dimensional forces and moments applied by the hands to the rim at 240 Hz, and has a precision of 0.6 N and resolution of 1 N [12].

During the comparison procedure, one able-bodied participant wore the glove while propelling a wheelchair instrumented with a SmartWheel along a concrete path with gentle turns and slopes for approximately ten minutes. The Mayo Clinic Institutional Review Board (IRB) determined this study was exempt from the requirement of IRB approval. Two weight relief exercises were performed at the beginning and end of the session.

**Table 1:** Loading conditions for all calibrations

| Condition | 1 | 2 | 3 | 4 | 5 |
| --- | --- | --- | --- | --- | --- |
| Substrate | Metal | Ballistic Gel | Synthetic Flesh | Synthetic Flesh | Ballistic Gel |
| Temperature (C) | 22 | 22 | 22 | 35 | 22 |
| Force Applicator | Flat | Flat | Flat | Flat | Wheelchair Rim |

During post-processing, the time-series force signals from the glove and SmartWheel were compared. Analyses were performed separately for weight relief and propulsion activities. The SmartWheel data were resampled from 240 Hz to 20 Hz, and the signals were aligned using the *alignsignals* Matlab function.

Propulsive peaks were counted in both the raw glove and SmartWheel signals using the *findpeaks* function. All peak indices were validated visually by an experimenter; mislabeled peaks were removed. Root mean square errors (RMSEs) in force were computed over the duration of the signals for both weight relief and propulsion. For every propulsion or weight relief exercise recorded by the glove, the percent error of the peak magnitude was computed as:

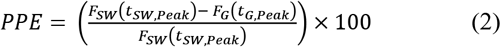

where PPE is Peak Percent Error, F_SW_(t) is the force magnitude measured by the SmartWheel at time t, F_G_(t) is the force magnitude predicted by the glove at time t, and t_SW,Peak_ and t_G,Peak_ are the times the SmartWheel and glove measured the peak, respectively. To compare across conditions, the mean absolute value for all PPEs were computed for weight relief and propulsion.

## III. Results

The results of the MTS calibration procedure are presented in Fig. 2 (top). The relationship between the glove and the MTS was generally linear (Table 2). However, the glove response varied substantially across calibration conditions for the same applied load, with condition one having the smallest range of response in the glove and condition five having the largest range of response.

**Figure 2:**
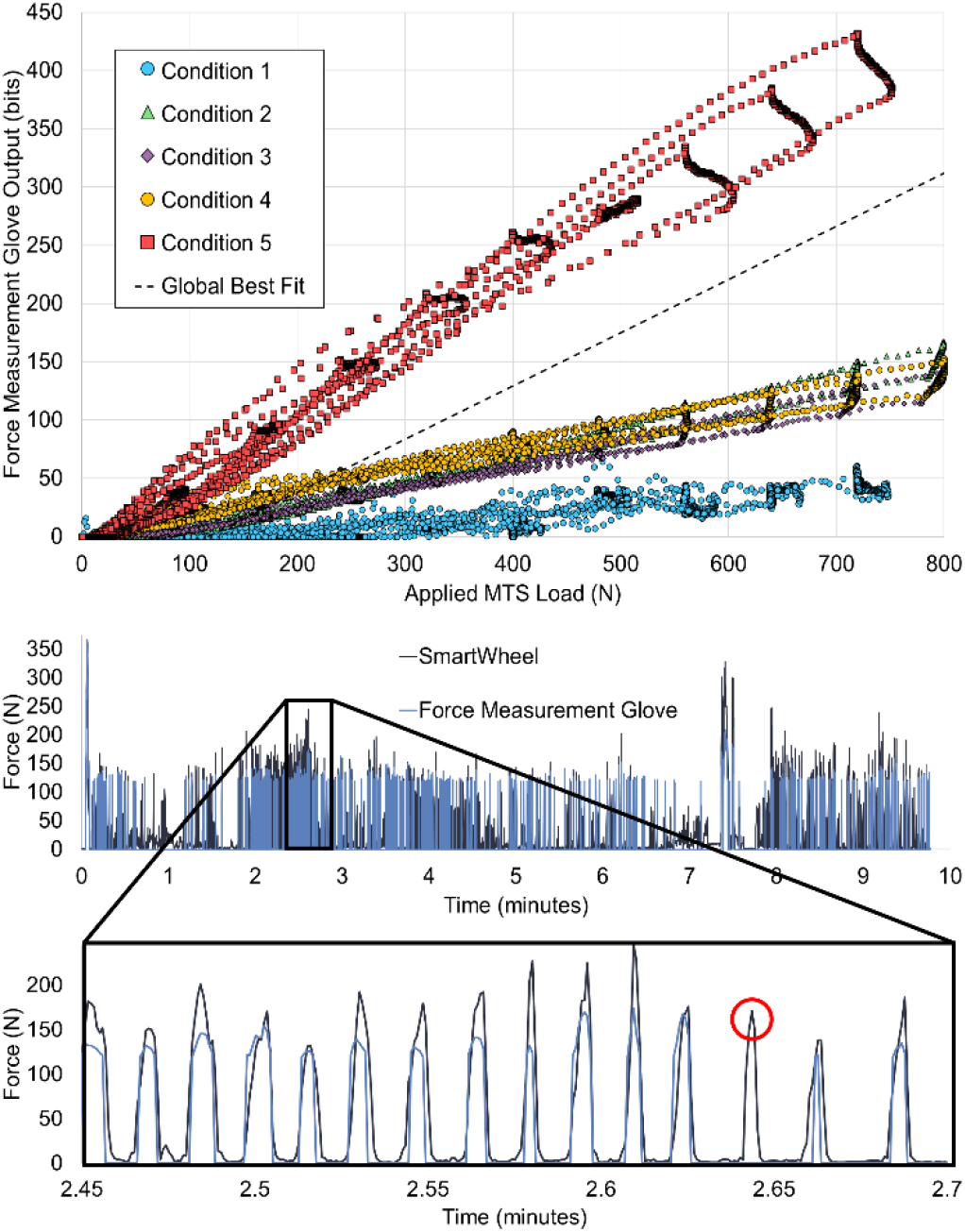
(Top) glove and MTS data from the five calibration conditions. This data was used to build six calibration equations. (Bottom) Time-series data from the SmartWheel validation procedure. (Bottom-inset) Subset of the force signal data during propulsion.

**Table 2:**
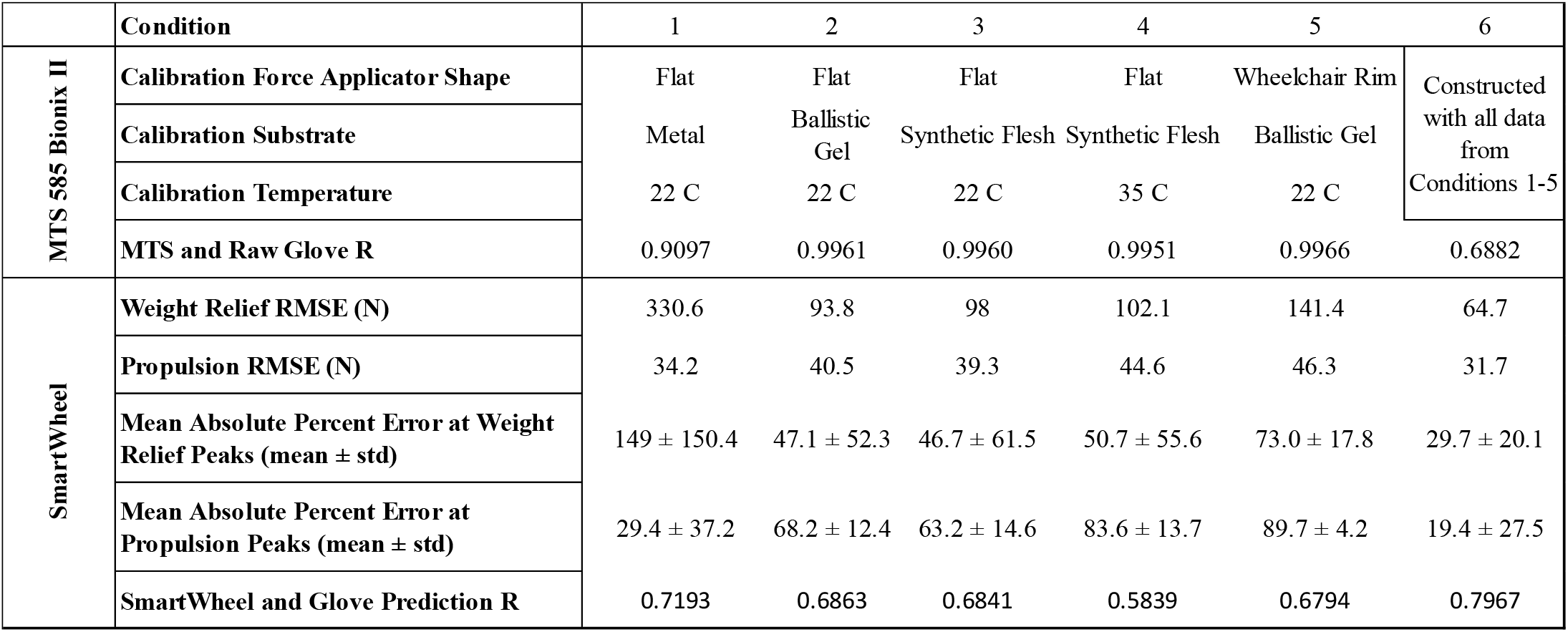
Error metrics for the calibration and SmartWheel comparison. Percent error calculations include 258 propulsion cycles and four weight relief exercises.

The raw glove signal contained 258 of the 355 peaks detected by the SmartWheel (72.7%). Fig. 2 (bottom-inset) shows the glove and SmartWheel signals during several propulsion cycles and an example of a propulsion cycle missed by the glove. The error metrics for the glove compared to the SmartWheel across conditions are detailed in Table 2. Calibration condition six, which consisted of all data from all calibration conditions, yielded the lowest error for all metrics and a linear correlation of r = 0.80. This implies that the average loading condition at the palm more closely aligned with the average of the five testing conditions than any of the of the individual conditions.

## IV. Discussion

In this study, we introduced an FSR-based force-measurement glove and investigated the effects of five different calibration conditions on measurement accuracy. We found that calibrating the glove with all data from a variety of biomechanically informed loading conditions minimized disagreement between glove force measurements and SmartWheel measurements and improved the linear correlation between the glove and SmartWheel measurements.

With an overall accuracy of wheelchair propulsion peak detection of 73% and a linear correlation of r = 0.80 between the glove and instrumented rim using calibration condition six, the glove showed moderate-to-low fidelity in detecting wheelchair propulsion loading events. The mean percent error for the magnitude of loading during the weight relief task using calibration condition six was 29.7% and the error was 19.4% for the wheelchair propulsion peak magnitudes. The missed loading peaks during propulsion were likely due to the participant’s hand placement and propulsion strategy. Forces generated at the fingers and parts of the palm did not pass through the FSRs in the glove; therefore, they were not measured. Within the glove design, the sensors on the palm are movable and perhaps the current placement was not optimal.

In comparison to a similar study [8], the sensors in the glove were designed for and calibrated at much higher loads (0-800 N vs. 0-10 N). For each loading condition, the glove exhibited linear input-output behavior whereas all sensors in Schofield 2016 exhibited exponential relationships between applied sensor output and load, which may be related to the specific FSR sensor type. The glove design has the capability to function with different sensors in individualized arrangements depending on the specific loading condition.

A major limitation of the study was the method of comparing the glove force prediction to the SmartWheel measurement, as the wheelchair rim measures a three-dimensional force vector while each sensor in the glove only measures force normal to its surface. The glove cannot measure shear forces at the palm, so we expect force measurements from the SmartWheel to be greater in magnitude than force predictions. An alternative approach to calibrate the glove with the SmartWheel data directly was not used due to the lack of commercial availability of the SmartWheel at the time of manuscript preparation.

Whether or not the measurement accuracies reported in this study for the force-measurement glove are good enough for extended field-based studies ultimately depends on the specific research question. The current glove can be used to count loading events over the course of a day, or detect high load events like body transfers, which have been implicated in shoulder pathology in manual wheelchair users. The current glove should not be used to measure absolute forces during manual wheelchair use due to the glove’s sensitivity to biomechanical variables at the palm.

Future work may focus on the development of a higher accuracy force measurement glove and optimizing sensor placement for wheelchair-based activities. A more advanced glove might include other types of sensors and machine learning methods to map glove measurements to force predictions and optimize sensor placement at the palm.

